# Evidence for *NR2F2*/COUP-TFII involvement in human testis development

**DOI:** 10.1101/2024.01.05.574424

**Authors:** Somboon Wankanit, Housna Zidoune, Joëlle Bignon-Topalovic, Laurène Schlick, Denis Houzelstein, Leila Fusée, Asma Boukri, Nassim Nouri, Ken McElreavey, Anu Bashamboo, Maëva Elzaiat

## Abstract

*NR2F2* encodes COUP-TFII, an orphan nuclear receptor involved in mammalian gonadogenesis. COUP-TFII is expressed in the interstitial/stromal compartment of both fetal testes and ovaries, where it is required for developing steroidogenic lineages. Pathogenic variants in human *NR2F2* are linked to testis formation in 46,XX individuals (46,XX <u>d</u>isorders of <u>s</u>ex <u>d</u>evelopment, 46,XX DSD). Such findings propose a regulatory role of COUP-TFII in the developing ovary, whereas its function in testis remains unknown. We evaluate the effect of a *de novo* heterozygous, predicted damaging, missense variant in *NR2F2* (p.Arg246His) in a 46,XY under-masculinized boy. *In-vitro* assays show that the mutant protein significantly loses the inhibitory effect on NR5A1-mediated activation of both the *LHB* and *INSL3* promoters. The data support the pathogenicity of the p.Arg246His variant in 46,XY DSD and a role for *NR2F2* in human testis formation. In addition to *NR5A1* and *WT1*, *NR2F2* variants are thus associated with both 46,XX and 46,XY DSD. This expands the list of genes that function in both male and female sex development, which is originally thought to be regulated by two entirely different sets of genes.

## Introduction

Sex determination is the process by which a sexually reproducing organism initiates differentiation as either a male or female. Commitment of a common progenitor to either male (Sertoli cell) or female (granulosa cell) fate is the outcome of a battle between poorly characterized organ specific, mutually antagonistic gene regulatory networks that canalize development down one organogenetic pathway, whilst actively repressing the alternate^1^. In XY individuals, *Sex determining region Y* (*Sry*) gene initiates a genetic cascade that leads to upregulation of *SRY-box transcription factor 9* (*Sox9*) beyond a critical threshold^1^.This results in the differentiation of bipotential somatic cell precursors into Sertoli cells that orchestrate testicular development. In the absence of SRY, in XX individuals, commitment to the granulosa cell lineage occurs via RSPO1/WNT4/β-CATENIN and RUNX1/FOXL2 gene regulatory networks^2,3^. Sex determination is followed by the differentiation of internal and external genitalia regulated by the presence or absence of androgens and anti-Müllerian hormone^4^.

Errors in sex determination/development cascades give rise to a heterogeneous group of pathologies termed disorders/differences of sex development (DSD). DSD are congenital conditions in which the development of chromosomal, gonadal or anatomical sex is discordant^5^. DSD can present in isolation or with multiple somatic anomalies in various syndromes^5^. Based on chromosomal composition, DSD are classified into three groups, including 46,XY DSD, 46,XX DSD, and sex chromosome DSD. 46,XY DSD encompass anomalies of testicular development, androgen synthesis oraction, persistent Müllerian duct syndrome and unclassified structural variations, and manifest as genital undervirilization^6^. 46,XX DSD includes testicular or ovotesticular DSD where XX individuals develop testis or ovotestis, and have virilized genitalia due to excessive androgen^7^. Despite recent technological advances, a definitive genetic diagnosis is achieved in less than 50% of 46,XY DSD cases and 20% of 46,XX testicular or ovotesticular DSD^8–10^. Etiologically, 46,XY and 46,XX DSD were considered to be distinct pathologies caused by mutations in different groups of genes, however, recent data provide evidence to the contrary. Pathogenic variants in *Nuclear receptor subfamily 5 group A member 1* (*NR5A1*) were initially associated with a wide range of reproductive phenotypes including 46,XY DSD, male infertility and primary ovarian insufficiency in 46,XX individuals^11–13^. Recently, a specific missense variant (p.Arg92) was described in associationwith 46,XX testicular or ovotesticular DSD^14–17^. Similarly, pathogenic variants of Wilms tumor 1 (*WT1*) gene were first reported in association with syndromic 46,XY DSD^18–21^, and subsequently variant impacting 4^th^ zinc finger of WT1 were described in individuals presenting with 46,XX testicular or ovotesticular DSD^22^.

*Nuclear receptor subfamily 2 group F member 2* (*NR2F2*) gene encodes the orphan nuclear receptor chicken ovalbumin upstream promoter-transcription factor II (COUP-TFII)^23^. COUP-TFII is involved in many vital processes such as metabolic homeostasis, angiogenesis, organogenesis, and cell fate determination and differentiation during embryonic development^24^. In the developing murine and human ovaries, COUP-TFII protein is detected in the interstitial/stromal population, presumed to be the precursor of theca cells^25,26^. In adult human ovaries, COUP-TFII co-localizes with the steroidogenic enzyme 17α-hydroxylase in the theca cells^27^. In fetal human testes, COUP-TFII is detected from 7 weeks of gestation in interstitial cells that later become Leydig cells^28^, where it physically interacts with NR5A1 to regulate *Insulin-like 3* (*Insl3*) gene expression required for testicular descent in mice^29–31^. COUP-TFII is also expressed in pituitary gonadotropes where it represses NR5A1-mediated activation of rat *Luteinizing hormone subunit beta* (*Lhb*) promoter^32^. Pathogenic variants in *NR2F2* have been associated with a range of pathologies, including facial dysmorphism, congenital heart defect, diaphragmatic hernia, asplenia and 46,XX DSD (OMIM#107773)^23,25,33–35^. To date, five patients have been reported with 46,XX testicular or ovotesticular DSD, together with developmental abnormalities of the eyelids, heart and diaphragm. These include two patients carrying *de novo NR2F2* frameshift variants (p.Gly35Argfs*75, p.Pro33Alafs*77), one with the p.Pro33Alafs*77 variant (unknown mode of transmission), one patient with a *de novo* missense variant (p.Trp8Ter), and another with a *de novo* deletion at 15q26.2 encompassing *NR2F2* locus^25,33,35^.

We previously reported the first case of non-syndromic 46,XY DSD involving a *de novo* heterozygous missense variant in *NR2F2*, NM_021005.4:c.737G>A causing p.Arg246His substitution^36^. The boy presented with micropenis, undescended testes and hypospadias. In this study, we evaluate the pathogenicity of the p.Arg246His variant. We found it did not impact nuclear localization, stability of the protein, nor the interaction with NR5A1 protein. However, we observed that COUP-TFII p.Arg246His significantly lost the inhibitory effect on NR5A1-dependent activation on target promoters including *LHB* and *INSL3*. As a result, our data support the pathogenicity of the *NR2F2* NM_021005.4:c.737G>A variant and expand the list of genes not only associated with 46,XX DSD, but also 46,XY DSD.

## Results

### Clinical phenotype of the patient

A 2-year-old 46,XY boy, of healthy non-consanguineous parents with North African ancestry, presented with under-virilized genitalia^36^. DSD was not reported in any other members of the family (Figure 1A). Clinical examination showed micropenis, middle hypospadias, well-formed scrotum, and inguinal testes. Müllerian structures were not observed on abdominopelvic ultrasonography while other visceral organs were unremarkable. Hormonal assessment at the presentation and reassessment at 5 years of age showed age-appropriate levels of serum gonadotropins, androgens and anti-Müllerian hormone (Table 1). There was no reported anomaly in other organ systems. The diagnosis was non-syndromic 46,XY DSD of unknown etiology.

**Figure 1.**
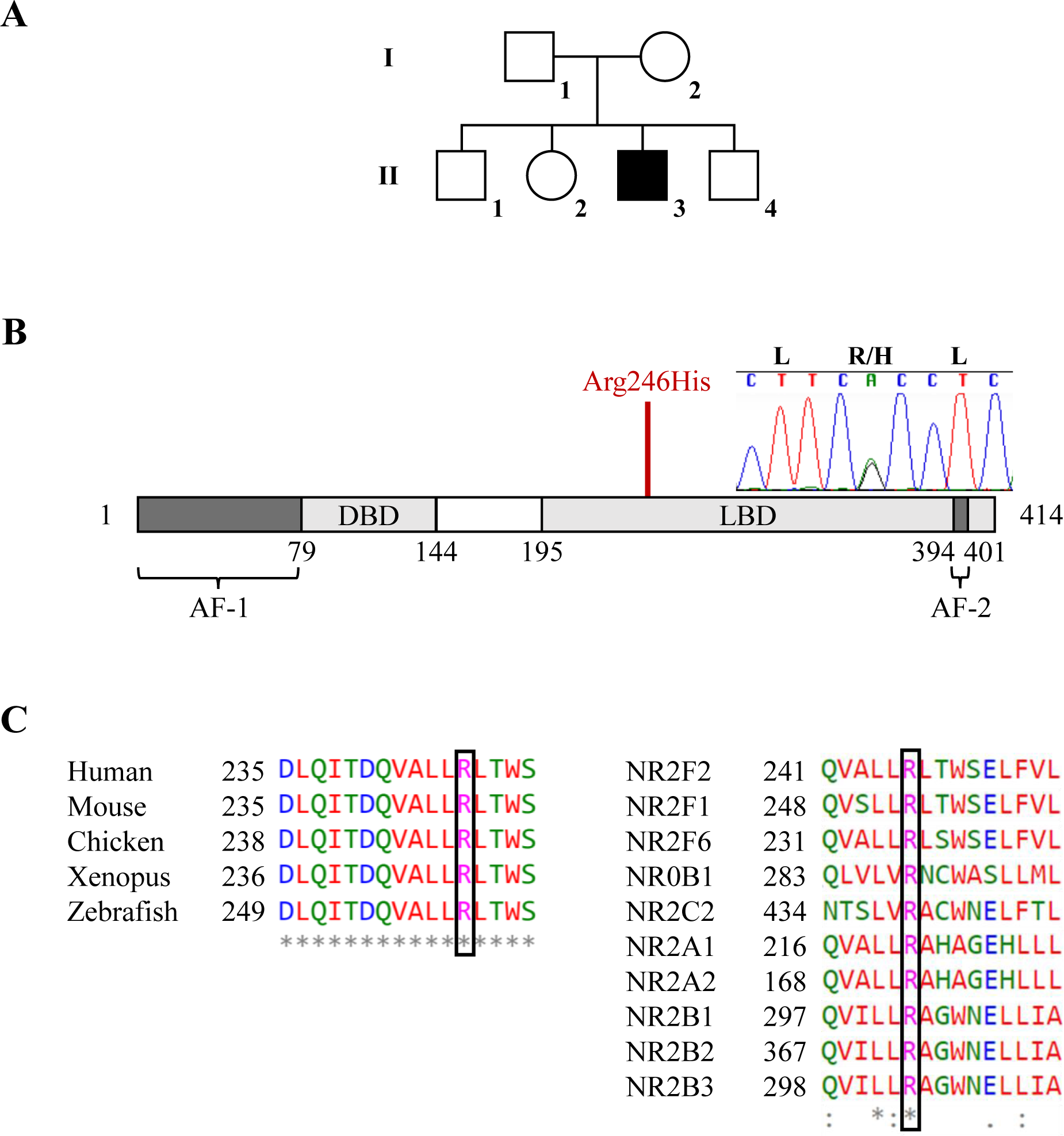
Identification and characteristics of the COUP-TFII p.Arg246His variant. (A) The pedigree of the family is indicated. Squares denote male family members and circles denote female family members. Solid square represents the patient. (B) Chromatogram shows the *de novo* heterozygous change of *NR2F2* (NM_021005: c.G737A > p.Arg246His). The functional domains of the COUP-TFII protein are shown and the mutated residue is located in the ligand-binding domain. (C) Comparative sequence alignment demonstrates that the variant falls in the highly conserved region across species (left) and in the ligand-binding domain of several nuclear receptors (right). AF, activator function; DBD, DNA-binding domain; LBD, ligand-binding domain.

**Table 1.** Hormonal profile of the patient.

| **Hormone** | **Age** |  | **Normal range** |
| --- | --- | --- | --- |
|  | **2 years** | **5 years** |  |
| Luteinizing hormone (IU/L) | 0.37 | <0.1 | <1.5 |
| Follicle-stimulating hormone (IU/L) | 1.22 | 0.53 | <5.0 |
| Testosterone (nmol/L) | <0.09 | 0.09 | <0.1-1 |
| hCG-stimulated testosterone (nmol/L) | 24 | - | $23 \pm 9^{59}$ |
| hCG-stimulated dihydrotestosterone (nmol/L) | 1.82 | - | $1.88 \pm 0.07^{59}$ |
| Anti-Müllerian hormone (pmol/L) | - | 291 | 90-1193 |

593 hCG, human chorionic gonadotropin

### The mutated residue is located within the ligand binding domain (LBD) and is predicted to affect protein function

Whole exome sequencing identified a novel heterozygous variant in *NR2F2* gene (NM_021005:c.G737A > p.Arg246His) (Figure 1B). Pathogenic variants in other known 46,XY DSD-causing genes were not identified^37^. Sanger sequencing of the patient and his parents confirmed that the *NR2F2* variant was *de novo*. The variant is absent in public databases (https://gnomad.broadinstitute.org/), is classified as likely pathogenic based on the American College of Medical Genetics criteria^38^, and is predicted to be deleterious by multiple algorithms including PolyPhen (0.988), SIFT (0.03), REVEL (0.922), MutationTaster (0.99) and CADD (32). The mutated residue is localized in the LBD of COUP-TFII (Figure 1B) and is highly conserved across species, as well as in the ligand-binding domain of several nuclear receptors (Figure 1C). HOPE software (https://www3.cmbi.umcn.nl/hope/)^39^ revealed that, unlike the positive charge of the wild-type (WT) residue arginine, the mutated histidine-residue charge is neutral,which is predicted to disturb the ionic interaction that the residue forms with the glycine at position 403, and consequently the protein conformation.

### The mutation does not impact the protein stability, subcellular localization nor interaction with NR5A1

We evaluated the effect of the mutation on the protein stability by Western blot. Both WT and mutant proteins were detected around 46 kDa and the quantity of both protein variants was comparable (*p*=0.84) (Figure 2A). We assessed if the mutant variant affected the subcellular localization of the resulting protein by immunocytochemistry. The mutant protein was localized in the nuclei, as observed in the WT protein (Figure 2B). The p.Arg246His mutated residue is located within the highly-conserved LBD, which plays a crucial role in the functions of nuclear receptors, including ligand recognition and ligand-dependent activation^40^. As COUP-TFII and NR5A1 are known to interact physically to regulate gene transcription^31^, we performed Duolink® proximity ligation assay to study the conservation in the binding to NR5A1 between the WT and mutant proteins. Quantification of the interactions between NR5A1 and WT or mutant COUP-TFII did not reveal significant difference (Figure 2C).

**Figure 2.**
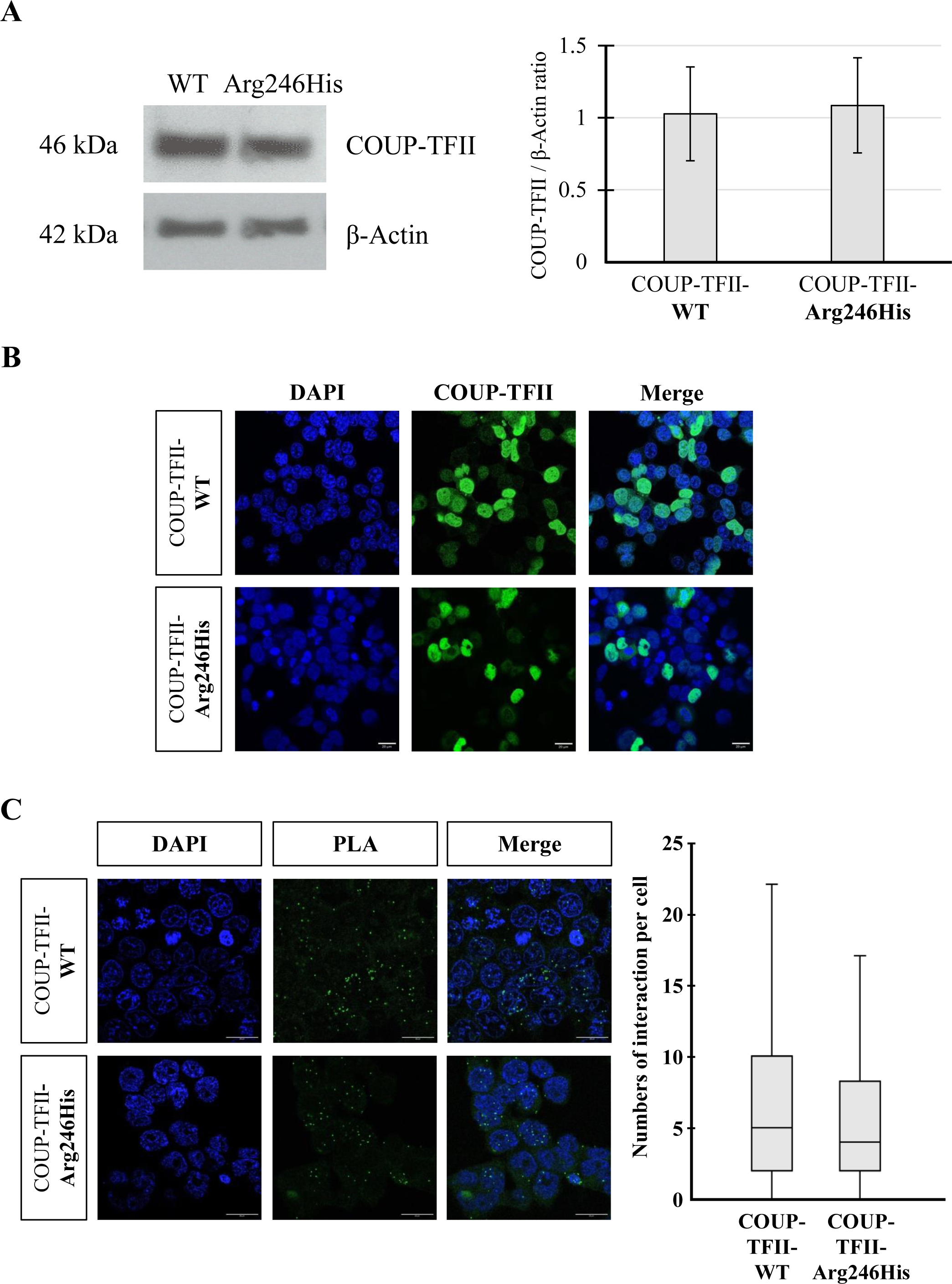
The mutation does not impact the variant stability, subcellular localization or interaction with NR5A1. (A) Protein stability. Left: protein extraction was performed after HEK-293T cells were transfected with either NR2F2-WT or NR2F2-p.Arg246His plasmids for 48 hours. A single band around 46 kDa was detected in both COUP-TFII-WT and COUP-TFII-p.Arg246His conditions. Right: the band intensity was normalized to the one of β-Actin and plotted as mean ± SEM of COUP-TFII/β-Actin ratio obtained from three independent experiments. There was no significant difference between the COUP-TFII-WT and COUP-TFII-p.Arg246His protein stability (Student t-test, *p* = 0.84). (B) Subcellular localization. HEK293-T cells were transfected with NR2F2-WT or NR2F2-p.Arg246His plasmids. The subcellular localization of the WT and mutant variants was analyzed by immunofluorescence. Both variants showed the same nuclear localization (green). Images were taken at 40X magnification (Scale bar: 20 µm). (C) The mutation has no significant impact on the interaction between COUP-TFII and NR5A1. Left: NR5A1 and NR2F2-WT or NR2F2-p.Arg246His plasmids were transiently expressed for 48 hours in HEK293-T cells. Protein-Protein interaction was assessed by the Duolink® proximity ligation assay (PLA). Nuclei are stained with DAPI (blue) while each PLA signal/dot (green) represents a COUP-TFII/NR5A1 interaction event. Right: quantification of PLA. For each condition, the number of interactions of at least 50 individual cells were counted and the median (interquartile range) was analyzed. Statistical analysis revealed that the binding of NR5A1 to COUP-TFII-p.Arg246His was comparable to that of COUP-TFII-WT (Mann-Whitney test, *p* = 0.22). Images were taken at 63X magnification (Scale bar: 20 µm).

### Mutant COUP-TFII alters the transactivation activity of NR5A1 on target promoters

We investigated whether the mutation altered the transcriptional activity of COUP-TFII. *LHB* and *INSL3* promoters are known targets of COUP-TFII and both of them are stimulated by NR5A1^31,32^. A previous study by Zheng *et al.* reported that COUP-TFII repressed NR5A1-dependent activation of rat *Lhb* promoter^32^. We assessed the transactivation ability of WT and mutant COUP-TFIIs using rat and human *Lhb*/*LHB* promoters in Human Embryonic Kidney 293 containing the SV40 large T antigen (HEK293-T) cells. We confirmed that WT COUP-TFII blunted the stimulatory effect of NR5A1 on rat *Lhb* promoter (Supplementary figure S1) while a similar result was observed on human *LHB* promoter (Figure 3A). The inhibitory effect was also significant when WT COUP-TFII was transfected alone, and may reflect endogenous NR5A1 transcriptional activity. The inhibitory effect of COUP-TFII p.Arg246His was significantly decreased (*p*<0.001). Notably, we observed comparable results when using human *INSL3* promoter (Figure 3B). WT COUP-TFII reduced the NR5A1-mediated activation on *INSL3* promoter while this effect was completely lost with the mutant COUP-TFII (*p*<0.001).

**Figure 3.**
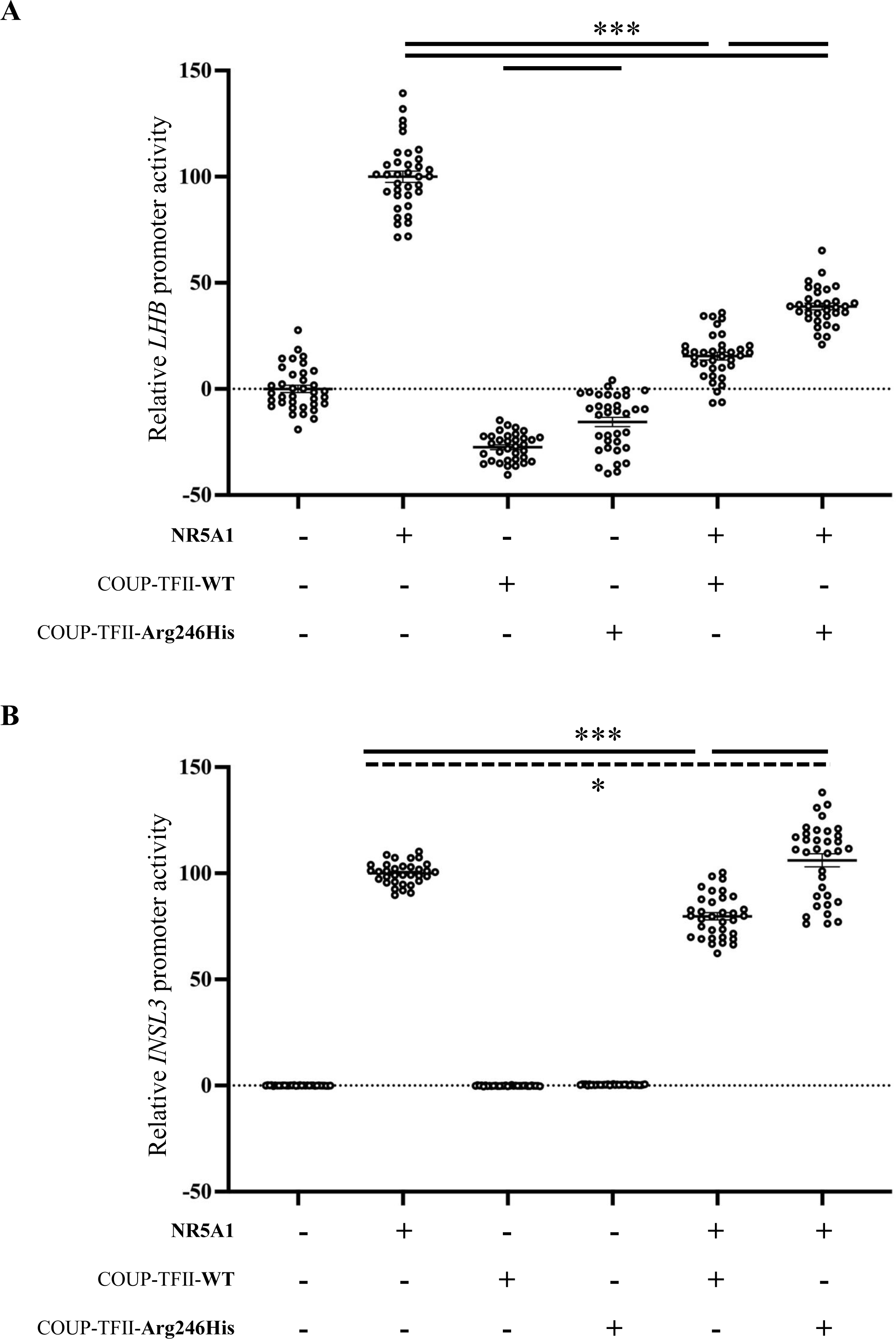
Mutant COUP-TFII shows loss of inhibitory effect on target promoters. The transcriptional activity of COUP-TFII-WT and COUP-TFII-p.Arg246His protein variants was assessed in presence or absence of NR5A1 on (A) *LHB* and (B) *INSL3* promoters fused to the *Luciferase* gene reporter. All data were standardized for Renilla activity. The data shown represent scatter dots of results with the mean ± SEM of three biological experiments, each with twelve replicates. The results are expressed in relative to the percentage of NR5A1 activation (100%). The statistical significances are marked by the horizontal lines (dash line*, *p*<0.05; bold line***, *p*<0.001).

## Discussion

We identified a heterozygous missense variant in *NR2F2* (NM_021005: c.G737A > p.Arg246His) in a male child presenting with non-syndromic 46,XY DSD. Predictive analysis supported the pathogenicity of the variant. COUP-TFII p.Arg246His did not affect the stability, subcellular localization, or the protein binding capacity with NR5A1. However, as compared to the wild type, the mutant protein failed to appropriately suppress NR5A1-dependent activation of target promoters.

The biological function of COUP-TFII in testis development has been inferred from mouse models, which indicate a key role for the protein in Leydig cell differentiation and androgen production via the regulation of genes encoding steroidogenic enzymes. The inducible depletion of *Nr2f2* at E18.5 in XY mice results in Leydig cell hypoplasia, whilst *Nr2f2*-depletion at pre-pubertal stages results in reduced expression of several genes encoding steroidogenic enzymes, including *Star*, *Cyp11a1*, *Hsd3b1* and *Cyp17a1*, as well as decreased testosterone biosynthesis^41^. In murine immortalized Leydig cell lines (MA-10 and MLTC-1), silencing COUP-TFII reduces expression of *Star*, which shuttles cholesterol from the outer to the inner mitochondrial membrane and initiates steroidogenesis, resulting in significantly decreased testosterone production^42^. This effect may also be caused by the reduced expression of *Glutathione S-Transferase Alpha 3* (*Gsta3*) that is required for the production of Δ4-androstene-3,17-dione, a precursor of testosterone^43^. In the human, COUP-TFII expression is detected from gestational week (GW) 7 in the interstitial compartment of fetal testes^28^. At GW8, Leydig cells can be identified in the interstitial tissue associated with initiation of testosterone production^28,44^. *NR2F2* expression is down-regulated in fetal Leydig cells by GW15 followed by a decrease in the testosterone levels^28,45^. Taken together these data indicate a regulatory role for human COUP-TFII in testosterone biosynthesis during the first trimester of gestation.

We hypothesize that the undervirilization of the external genitalia observed in the individual described in this study could be caused by reduced prenatal testosterone levels either due to hypogonadotropic hypogonadism or primary hypogonadism. In humans, testosterone production is regulated by placental human chorionic gonadotropin during the first trimester of gestation, and subsequently by luteinizing hormone of the fetal pituitary from mid-gestation^46^. Testosterone is required for the formation of male external genitalia within the first trimester, and partly for penile growth and inguinoscrotal descent of testes during the third trimester^46^. Therefore, anomalies of external genitalia, including micropenis, cryptorchidism and hypospadias are common in patients with 46,XY DSD, in whom testosterone synthesis or action is impaired from an early gestational stage^47^. In contrast, hypospadias is rarely observed in patients with congenital hypogonadotropic hypogonadism where luteinizing hormone secretion is low and may impact genital development at a later stage^46,48,49^. In addition, less than 30% of patients with congenital hypogonadotropic hypogonadism present with micropenis and undescended testes^49^. Therefore, genital undervirilization including micropenis, undescended testes together with hypospadias support the diagnosis of 46,XY DSD due to primary hypogonadism.

We characterize the first *NR2F2* variant associated with non-syndromic 46,XY DSD. Genetic abnormalities involving the *Nr2f2/NR2F2* gene in both mouse and human males were previously associated with genital undervirilization associated with somatic anomalies^34,50–55^. In mice, the conditional deletion of *Nr2f2* causes cryptorchidism in addition to cardiovascular and diaphragmatic defects^52^. Chromosomal aberrations encompassing the 15q26.2 region, the locus for human *NR2F2*, were reported in boys with multiple dysmorphic features (Supplementary Table 1)^50,51,53–55^. These chromosomal rearrangements ranged from 36 kb to 8.6 Mb in size^50,51,53–55^, and involved approximately 60 genes^54^. The genital anomalies included undescended testes, hypospadias, micropenis and scrotal hypoplasia, however, due to the size of 15q26.2 deletions, it was not possible to directly corelate these anomalies with *NR2F2*. A variant located in the LBD of *NR2F2*, p.Ser341Tyr, was reported in a 7 year-old boy with multiple congenital anomalies, including facial dysmorphism, microcephaly, mild global developmental delay, asplenia, glandular hypospadias and cryptorchidism^34^. However, it is unclear if the DSD in this patient is due to the p.Ser341Tyr variant or secondary to pituitary anomalies associated with the microcephaly as clinical and hormonal data at puberty is unavailable because of the patient’s young age at the presentation^34^.

Based on crystal structure, COUP-TFII is in an auto-repressed conformation in the absence of ligands^40^. The interaction between the activation-function 2 (AF-2) of the LBD and the co-factor binding site stabilizes the structure and prevents the recruitment of co-activator or co-repressor.Therefore, the AF-2 conformational states predominately determine the activity of nuclear receptors^56^. The AF-2 of COUP-TFII is stabilized by a hydrogen bond between Arg 246 and Glu 393 amino acid residues^40^. *In-vitro* studies show mutagenesis of residues 249 and 250 which lie adjacent to Arg 246 and line the LBD pocket caused a 50% decrease in transactivation of target *Nerve Growth Factor-Induced protein A* (*NGFI-A*) promoter compared to the WT protein^40^. This suggests that the COUP-TFII p.Arg246His variant disrupts the conformation of AF-2 domain, and consequently alters the transcriptional activity of the protein. This is supported by *in-vitro* assays in the present study, that indicate a significant loss of function for COUP-TFII p.Arg246His. In pituitary gonadotropes, COUP-TFII acts as a transcriptional repressor of NR5A1 mediated activation of rat *Lhb* promoter^32^. Of the two NR5A1-binding elements identified on rat *Lhb* promoter, COUP-TFII binds to the 3’-domain in competition with NR5A1 and thereby decreases NR5A1-mediated activation^32^. Unlike the 5’-binding element, the sequence of the 3’-binding element is conserved in human *LHB* promoter (Supplementary Figure S2). We observed that COUP-TFII decreased NR5A1-mediated activation of human *LHB* promoter similar to that of rat *Lhb* promoter (Figure 3A). However, the COUP-TFII p.Arg246His variant results in a statistically significant loss of this inhibitory effect on human *LHB* promoter as compared to the WT protein. Moreover, previous studies have shown that COUP-TFII acts synergistically with NR5A1 to enhance *INSL3* promoter activity in mouse Leydig cells and CV-1 fibroblast cells^31,57^.This synergy is proposed to be, in part, due to the recruitment of another transcription factor(s) acting as a co-activator^31^.We demonstrated that COUP-TFII decreases the NR5A1-dependent activation of *INSL3* promoter in HEK293-T cells, while the COUP-TFII p.Arg246His completely lost this repressing effect. Although COUP-TFII is primarily known as a transcriptional repressor, it can also function as a transcriptional activator depending on the cellular and genetic context^23^. Therefore, in our study, the disparate observation of COUP-TFII on NR5A1-dependent *INSL3* promoter could be due to the use of different cell lines that have distinct endogenous transcription factors and cofactors.

Apart from *NR5A1* and *WT1* variants that are associated with either 46,XX or 46,XY DSD^11,18,20,22,25^, our data add *NR2F2* to this small but growing gene list. Such pathological findings provide a better understanding on factors involved in human sex development. Although different sets of gene are known to regulate male and female pathways, overlapping genes between the two cascades exist. Furthermore, the fact that none of the variants in *NR5A1*, *WT1* and *NR2F2* causes both 46,XX and 46,XY DSD indicates distinct functions of these genes between testis and ovary, and/or sex-specific partners that remain to be identified.

## Materials and methods

### Patient samples

The patient met the revised 46,XY criteria of the Pediatric Endocrine Society (LWPES)/European Society for Pediatric Endocrinology (ESPE). The study was approved by the local French ethical committee (2014/ 18NICB-registration number IRB00003835). Family history including the birthplace, language, and ethnicity of the participant and his parents was obtained by self-reporting.

### Whole exome sequencing

Following whole exome sequencing, exon enrichment was performed using Agilent SureSelect Human All Exon V4. Paired-end sequencing with an average sequencing coverage of ×50 was performed on the Illumina HiSeq2000 platform. The sequencing platform of the manufacturer’s proprietary software was applied for generating read files which were then mapped with the Burrows-Wheeler Aligner to the human genome reference (hg38, http://hgdownload.cse.ucsc.edu/goldenPath/hg38/bigZips/analysisSet/hg38.analysisSet.2bit). Duplicate reads were marked by using Picard. Additional BAM files were sorted by SAMtools. The GATK version 1.6 was used for local realignment of the mapped reads around potential insertion/deletion (indel) sites. SNP and indel variants of each sample were called by the GATK Unified Genotyper. SNP novelty was determined against dbSNP138. The fractions of exome targets with >20× and >10× coverage were>92% and >98%, respectively. We filtered out all known common variants that existed in dbSNP (build 138) (www.ncbi.nlm.nih.gov/projects/SNP/), the 1000 Genomes Project (http://www.1000genomes.org/), and the gnomAD database (http://gnomad.broadinstitute.org/). Variants with minor allele frequency (MAF) of >0.01 in any specific subpopulation were also excluded. After dataset filtering, novel or rare (MAF <0.01) variants were detected and analyzed by using the EnsEMBL SNP Effect Predictor (http://www.ensembl.org/homosapiens/userdata/uploadvariations). We focused our analyses on non-synonymous coding, nonsense, and splice-site variants. Manual screening against the Human Gene Mutation Database Professional (Biobase) (http://www.biobase-international.com/product/hgmd) was carried out for all variants. The remaining novel or rare variants were confirmed by visual examination using the IGV browser. The potential pathogenicity of the variants was determined by *in silico* analysis and potential pathogenic variants were confirmed by Sanger sequencing.

### In silico analysis

Classification of variants identified in this study was performed by using current population databases online tools including GnomADv3.1.2 (gnomad.broadinstitute.org) for analysis of allele frequency, and Ensembl genome browser 108 (https://www.ensembl.org/index.html) for predicting pathogenicity. Predictions on the structural and functional effects of the variant on the corresponding COUP-TFII protein (UniProt sequence #P24468) were assessed using HOPE software (http://www3.cmbi.ru.nl/hope/)^39^.

### Plasmids and Site-directed mutagenesis

pCMV6-XL5 NR2F2 (pCMV6-NR2F2, #SC108069, Origene) was used for evaluating COUP-TFII functions in this study. *NR2F2* expression vector containing the NM_021005.4:c.737G>A (p.Arg246His) variant was generated by site-directed mutagenesis (QuikChange II, Stratagene), using the forward primer: 5’-GACCAGGTGGCCCTGCTTCACCTCACCTGGAGCGAGC-3’ and the reverse primer 5’-GCTCGCTCCAGGTGAGGTGAAGCAGGGCCACCTGGTC-3’. The entire coding region sequence of the mutant plasmid was confirmed by direct Sanger sequencing. The pCMX-NR5A1 vector containing NR5A1 WT cDNA sequence has been described elsewhere^58^. The pCMX(https://www.addgene.org/vector-database/2249/) and pCDNA6 (#V22120, ThermoFisher Scientific) empty vectors were used as controls, as previously described^12^. The Luciferase reporter plasmid containing *INSL3* promoter was a generous gift from Prof. Jacques J. Tremblay and described in a previous study^43^. The *Lhb* promoter sequences from the rat (−207/+5)^32^, and the equivalent region in the human (−207/+5) were cloned into pGL3-basic Luciferase reporter-vector (E1751, Promega) using the forward primer containing XhoI restriction site: 5’-CGGGCTCGAGTTCCCAATGTCAGTTAAGC-3’ and the reverse primer containing SacI restriction site: 5’-GGTACCGAGCTCTTCCCAATGTCAGTTAAGC-3’ for the rat sequence, and the forward primer containing XhoI restriction site: 5’-CGGGCTCGAGGTCTCTGCCTCACCTCT-3’ and the reverse primer containing SacI: 5’-GGTACCGAGCTCGTCTCTGCCTCACCTCT-3’ for the human sequence. Plasmids were amplified following heat-shock transformation of NEB-5alpha competent cells (#C2992H, BioLabs), and purified with the NucleoBondXtra Maxi Plus kit (#740416, Macherey-Nagel).

### Cell culture

HEK293-T cells were cultured in DMEM medium (#31966-021, Gibco) supplemented with 10% fetal bovine serum (#10270-106, Gibco) and 1% Penicillin/Streptomycin 10,000 U/mL (#P06-07100, PAN Biotech). The cells were passaged every two days when cellular confluence reached 70-90%.

### Transient gene expression assays

For each well of 96-well plates (#0030730119, Eppendorf), 25,000 HEK293-T cells were seeded and transfected with FuGENE 6 (#E231A, Promega) and plasmid DNA at the carrier (µL):DNA (µg) ratio of 3:1. The transfection protocol was derived from previous studies on *LHB* and *INSL3* promoters^32,43^. Optimal transfection conditions were tested (Supplementary figure S3). Then, 50 ng of *LHB* promoter-containing reporter vector were transfected with 5 ng of pRLRenilla (#E2231, Promega), with or without 50 ng of pCMV6-NR2F2-WT or p.Arg246His and/or 5 ng of pCMX-NR5A1. A hundred ng of *INSL3* promoter-containing reporter vector was transfected with 5 ng of pRLRenilla, with or without 6.25 ng of pCMV6-NR2F2-WT or p.Arg246His and/or 6.25 ng of pCMX-NR5A1. Cells were lysed 48 hours after transfection. The promoter activity was measured based on the production of luminescence by using the Dual-Luciferase Reporter Assay system (#E1910, Promega) and the Glomax Multi+ Detection System (Promega). All data were standardized for Renilla activity. Results were plotted as the mean ± standard error of the mean (SEM) of three independent experiments, each with 12 replicates. Statistics were performed using RStudio. Outliers were determined by the interquartile range method, and removed. Pairwise comparisons were performed using the approximative two-sample Fisher-Pitman permutation test with stratification of the three independent experiments (Rstudio, coin package, one way-test). A *P*-value of less than 0.05 was considered as statistically significant.

### Immunofluorescence

To study the subcellular localization of the COUP-TFII variants, 20,000 HEK293-T cells were seeded in each well of LabTek chamberslides (#055071, Dutscher) and were transfected using FuGENE 6 with 100 ng of either pCMV6-NR2F2-WT or p.Arg246His. After 48 hours, the medium was removed. Cells were rinsed with phosphate-buffered saline (PBS) (#D8537, SIGMA) and were then fixed with 4% paraformaldehyde (PAF) for 10 minutes. After three washes with PBS, cells were permeabilized with PBS-0.1% Triton X-100 (#0694-1L, Amresco), and saturated with PBS-5% Bovine Serum Albumin (BSA) (#A3059-50G, SIGMA), each for 30 minutes. Cells were incubated overnight with rabbit anti-NR2F2 antibody (1:200, #ab42672, Abcam) diluted in PBS-3% BSA at 4°C. The following day, cells were washed thrice and incubated for 45 minutes with secondary antibody: Alexa488 goat anti-rabbit (1:1,000, #ab205718, Abcam) diluted in PBS-3% BSA. After counterstaining with 4′,6-diamidino-2-phenylindole (DAPI, 1:5,000, #62248, Thermo Scientific), slides were mounted using ProLong Gold Antifade Mountant with DAPI (#P39941, ThermoFisher Scientific). Images were obtained using Zeiss LSM900 confocal microscope at 40X magnification and analyzed using ImageJ.

### Duolink® proximity ligation assay (PLA)

To study the interaction of the NR2F2 variants with NR5A1, 20,000 HEK-293T cells were seeded in each well of a LabTek chamberslide (#055071, Nunc). Cells were co-transfected using FuGENE 6 with 100 ng of pCMV6-NR2F2-WT or p.Arg246His and 100 ng of pCMX-NR5A1. After 48 hours of culture, cells were fixed as described in the immunofluorescence section. The Duolink PLA was performed following the manufacturer’s instructions. In brief, cells were washed with PBS and permeabilized with PBS-0.1% Triton X-100 for 30 minutes. They were then incubated with the blocking solution (supplied in the kit) for an hour. Subsequently, overnight incubation at 4°C with both mouse anti-NR5A1 (1:200, #sc-393592, Santa-Cruz Biotechnologies) and rabbit anti-NR2F2 (1:200, #ab42672, Abcam) primary antibodies was performed. On the next day, samples were rinsed and incubated successively with PLUS (anti-Rabbit) (#DUO92002-100RXN, SIGMA) and MINUS (anti-Mouse) (#DUO92004-100RXN, SIGMA) PLA probes, ligation solution and amplification solution (#DUO92014-100RXN (Green), SIGMA) at 37°C. Following several washes, slides were mounted with Duolink In situ Mounting Medium with DAPI (#DUO082040, SIGMA). Finally, images were obtained by using Zeiss LSM900 confocal microscope at 63X magnification and analyzed with ImageJ software. For each condition, the number of interactions were counted in at least 50 individual cells and the median (interquartile range) of interaction numbers was compared using Mann-Whitney test. A *p*-value of less than 0.05 was considered as statistically significant.

### Protein preparation

A total of 350,000 HEK293-T cells were seeded in each well of 6-well plates (#EP0030720113, Eppendorf). Transfection was performed the day after at 40-50% cell confluence, using 3 µL of FuGENE 6 and 1 µg of pCMV6-NR2F2-WT or p.Arg246His in each well. After 48 hours of culture, cells were rinsed with PBS and lysed with IP lysis buffer (#87788, ThermoFisher Scientific) supplemented with Halt Protease Inhibitor Cocktail 100X (#78440, ThermoFisher Scientific) and EDTA 100X (#78440, ThermoFisher Scientific) for 30 minutes. The lysates were centrifuged for 15 minutes at 20,000g and the supernatants were retrieved. Protein quantification was performed using the Pierce^TM^ Detergent Compatible Bradford Assay kit (#23246, ThermoFisher Scientific) according to the manufacturer’s instructions.

### Western blot

For protein denaturation, 10 µg of protein samples was incubated with 4X XT loading buffer (#1610791, Bio-Rad) at 95°C for 5 minutes. Proteins were then separated on Criterion XT 10% polyacrylamide gel (#3450112, Bio-Rad) and transferred to PVDF membrane (#T831.1, Merk Millipore). Membranes were blocked in Tris-buffered saline containing 0.1% TWEEN 20 (#27949, SIGMA) (TBS-T) and 5% non-fat powdered milk for an hour at room temperature. After overnight incubation at 4°C with rabbit anti-NR2F2 (1:2000, #ab42672, Abcam) diluted in TBS-T-5% BSA, membranes were washed thrice with TBS-T. Then, membranes were incubated with anti-rabbit (#ab205718, Abcam) IgG antibody coupled to the Horse Radish Peroxidase (HRP) for 45 minutes. The revelation was performed using Pierce ECL Western blotting substrate (#32132, ThermoFisher Scientific) and X-Ray films. The detection of several proteins on the same blot was achieved by using the Antibody Stripping solution (#L7710A, Interchim) followed by probing with new primary antibodies. β-Actin detection was used for normalization. Band intensity was quantified using ImageJ software, and results were plotted as the mean ± SEM of three independent experiments using Excel. The statistical comparison of the means was performed using the student t-test. A *p*-value of less than 0.05 was considered as statistically significant.

## Acknowledgements

We are grateful to the patient and his parents for their participation in this study. We thank Prof. Jacques J. Tremblay for providing the Luciferase reporter plasmid containing human *INSL3* promoter.

## Ethics declarations

This study was approved by the local French ethical committee (2014/18NICB; registration no. IRB00003835) and Independent Ethical Committee at Hospital de Pediatria Garrahan (2016/971). Consent to genetic testing was obtained from the parents for this study.

## Author contributions

SW, HZ, KM, ABa and ME contributed to the conception and design of the study. HZ, ABo and NNprovided samples and detailed clinical information for the patient. SW, HZ, DH, LF, KM, ABa and ME performed the analysis of the datasets and wrote the manuscript. JBT and LS provided technical services. All authors contributed to manuscript revision, read, and approved the submitted version.

## Data availability statement

The data presented in the study are deposited in the ClinVar repository (https://www.ncbi.nlm.nih.gov/clinvar/).

## Supplementary Methods

### Transfection protocol for the transactivation assay on rat *Lhb* promoter

HEK293-T cells were seeded at 25,000 cells per well of 96-well plates (Eppendorf), and transfected with FuGENE 6 and plasmid DNA at the carrier (µL):DNA (µg) ratio of 3:1. Plasmid DNA including 60 ng of rat *Lhb* promoter fused to Luciferase reporter, 15 ng of pCMV6-NR2F2 WT or p.Arg246His, 15 ng of pCMX-NR5A1 and 5 ng of pRL Renilla was co-transfected into the cells which were lysed at approximately the next 48 hours.

### Condition testing for transactivation assay on human *LHB* and *INSL3* promoters

25,000 HEK293-T cells were seeded on each well of 96-well plates (Eppendorf), and transfected with FuGENE 6 and plasmid DNA at the carrier (µL):DNA (µg) ratio of 3:1. The amount of plasmid DNA for transactivation on human *LHB* promoter (p*LHB*) is shown in the table.

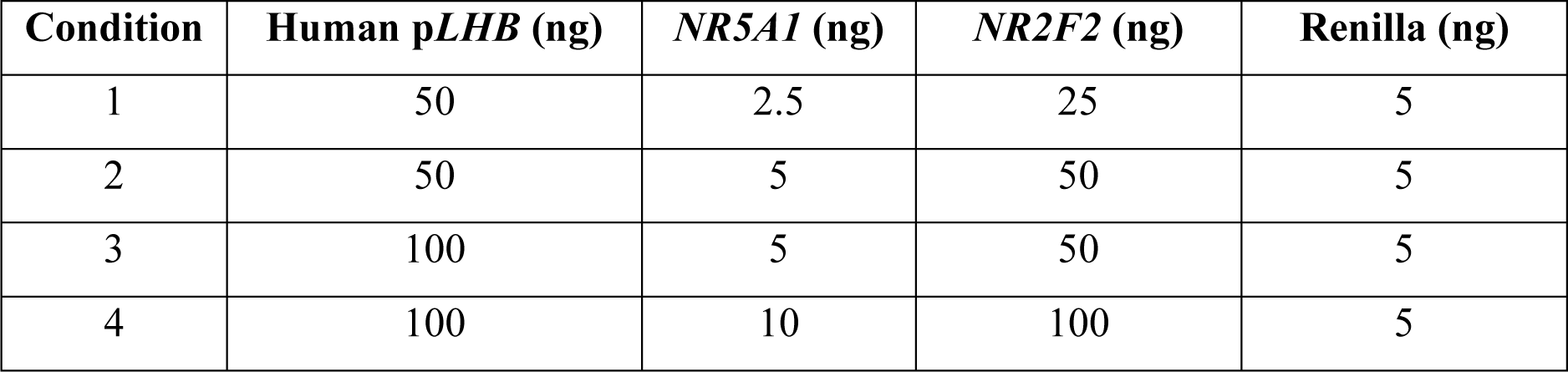

For the transactivation on human *INSL3* promoter (p*INSL3*), the amount of plasmid DNA tested was as follows.

| Condition | Human p*INSL3* (ng) | *NR5A1* (ng) | *NR2F2* (ng) | Renilla (ng) |
| --- | --- | --- | --- | --- |
| 1 | 50 | 3.1 | 3.1 | 5 |
| 2 | 50 | 6.2 | 6.2 | 5 |
| 3 | 100 | 6.2 | 6.2 | 5 |
| 4 | 100 | 12.5 | 12.5 | 5 |

## Supplementary figure legends

Supplementary figure S1. **COUP-TFII-p.Arg246His shows loss of inhibitory effect on rat *Lhb* promoter**. The transcriptional activity of COUP-TFII-WT and COUP-TFII-p.Arg246His protein variants was assessed in the presence of NR5A1 on rat *Lhb* promoter fused to the Luciferase gene reporter. All data were standardized for Renilla activity. The data shown represent the mean ± SEM of six replicates. The results are expressed in relative to the percentage of NR5A1 activation (100%).

Supplementary figure S2. **Comparative alignment of 5’-flanking sequences of rat and human *Lhb/LHB* promoters.** 5’ and 3’ NR5A1-binding elements of rat *Lhb* promoter are shown in bold. The conserved sequence of 3’ NR5A1-binding domain of human *LHB* promoter to which COUP-TFII competes with NR5A1 for the binding and represses NR5A1-dependent activation is highlighted in grey.

Supplementary figure S3. **Optimizing the conditions of Luciferase assay**.The transcriptional activity of NR5A1 and COUP-TFII-WT proteins was assessed on (A) *LHB* and (B) *INSL3* promoters fused to the Luciferase gene reporter. Different DNA quantities and ratios were tested. All data were standardized for Renilla activity. The data shown represent the mean six replicates. The results are expressed in relative to the percentage of NR5A1 activation (100%).

**Supplementary table 1.** Reports of boys with genital undervirilization and chromosomal aberration involving the 15q26.2 locus.

| Reference | Karyotype | Deletion size | Genital phenotype |
| --- | --- | --- | --- |
| Rump, *et al.* | 15q26.2-qter deletion | 5.8 Mb | Cryptorchidism, hypospadias, micropenis and hypoplastic scrotum |
| Choi, *et al.* | 15q26.2-qter deletion | 8.58 Mb | Cryptorchidism |
| Baptista, *et al.* | 46,XY,t(14;15)(q23;q26.3) | 36 kb | Cryptorchidism |
| Klaassens, *et al.* | 46,XY,t(1;14),inv(6),del(15)(q26) | 6.3 Mb | Genital anomalies |
| Butler, *et al.* | 46,XY,r(15)(p11q26) | Not available | Cryptorchidism and hypospadias |

