## Supplementary Figures 1-3 for "Evidence for *NR2F2*/COUP-TFII involvement in human testis development"

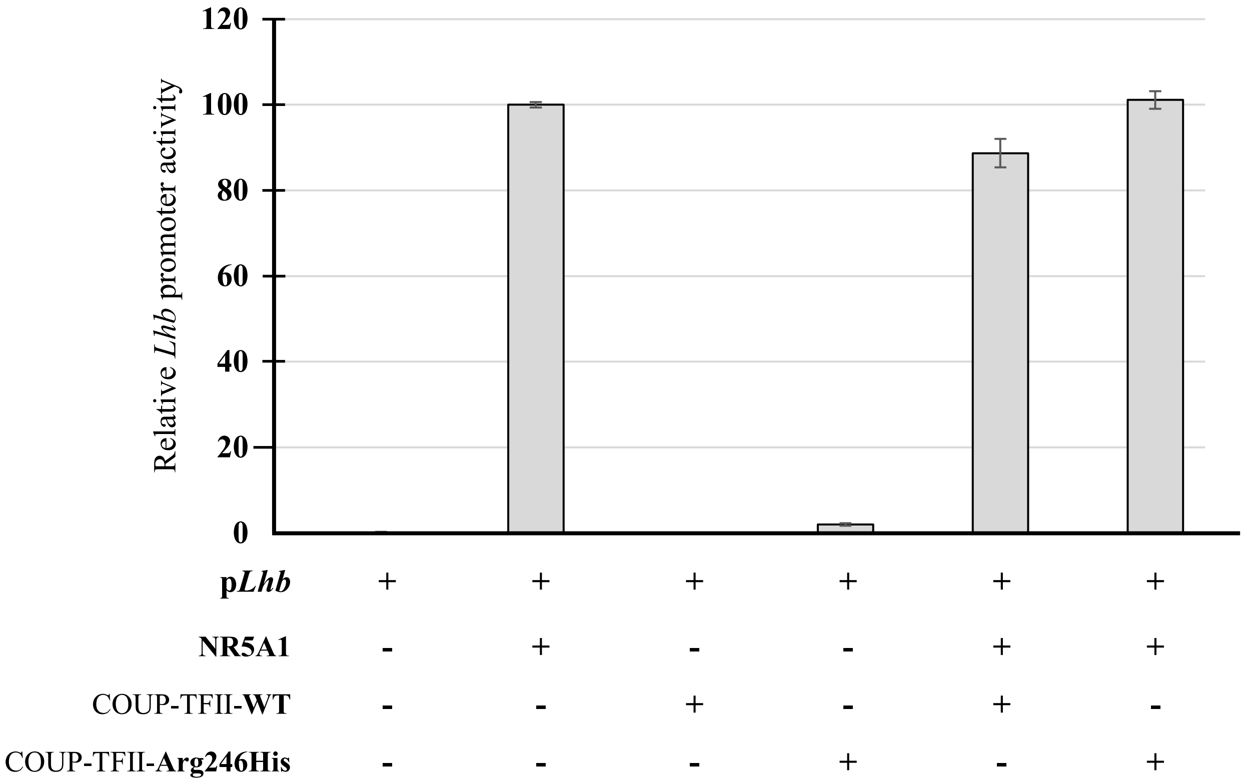


Supplementary Figure S1.

```
Rat        ---TTCCCAATGTCAGT--TAAGCTCAGGCACCT--GGGCTGAGTGTGAGGCCAATTCAC   53
Human      GTGGTCTCTGCCTCACCTCTGGCGCTAGACCACTGAGGGGAGAGGGCTGGGGCGCTCTGC   60
               ** *    ***    *      ** *  **  ***  *** *   ** *  *   *

Rat        TGAGACACTGGAGCTGGTCCCTGGCTTTTCTGACCTTGTCTGTCTCGCCCCCAAAGAGAT   113
Human      TGAGCCACTCCTGCGCCTCCCT-------GGCCATGTGCACCTCTCGCCCCCGGGGGGAT   113
           **** ****   **   *****              **    **********   * ***

Rat        TAGTGTCTAGGTTACCCAAGCCTGTAGCCTCTGCTTAGTGGCCTTGCCACCCCCACAACC   173
Human      TAGTGTCCAGGTTACCCCAGCATCCTATCACCTCCTGGTGGCCTTGCCGCCCCCACAACC   173
           ******* ********* *** *     * *  * * *********** ***********

Rat        CGCAGGTATAAAGCCAGGTGCCCAAGGTAGGGAAGGTAT 212
Human      CCGAGGTATATAGCCAGATACACGAGGCAGGGGATGCAC 212
           *  ******* ****** * * * *** **** * * *
```

Supplementary Figure S2.

**A**

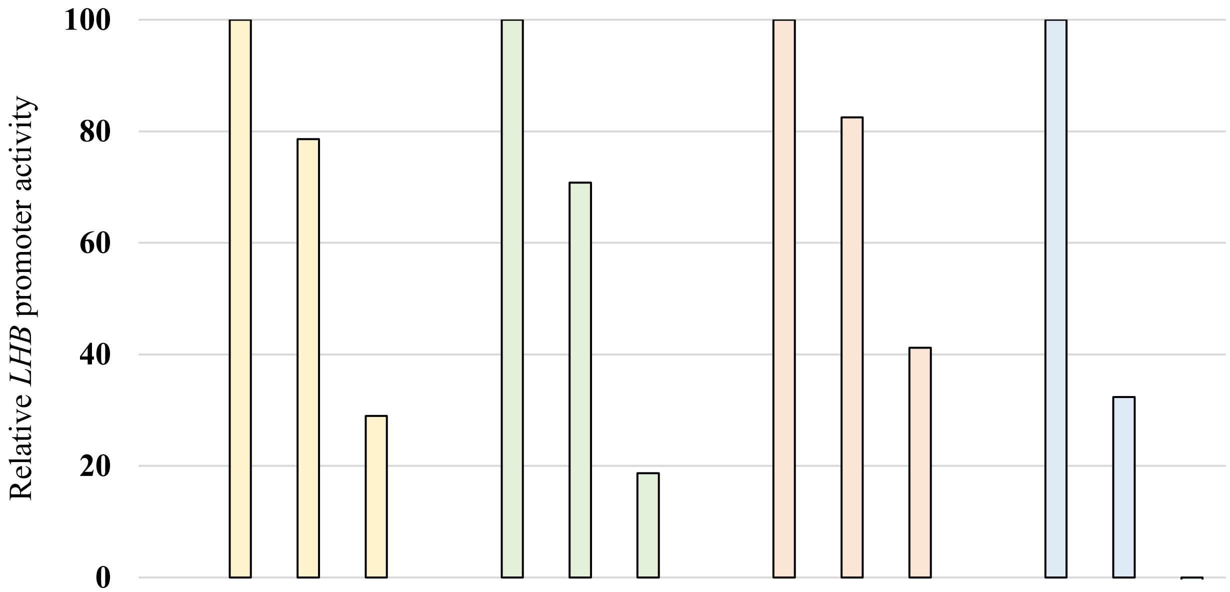


| | | | | | | | | | | | | | | | | |
|---|---|---|---|---|---|---|---|---|---|---|---|---|---|---|---|---|
| **p*LHB* (ng)** | 50 | 50 | 50 | 50 | 50 | 50 | 50 | 50 | 100 | 100 | 100 | 100 | 100 | 100 | 100 | 100 |
| **NR5A1 (ng)** | - | 2.5 | 2.5 | 2.5 | - | 5 | 5 | 5 | - | 5 | 5 | 5 | - | 10 | 10 | 10 |
| **COUP-TFII-WT (ng)** | - | - | 2.5 | 25 | - | - | 5 | 50 | - | - | 5 | 50 | - | - | 10 | 100 |

**B**

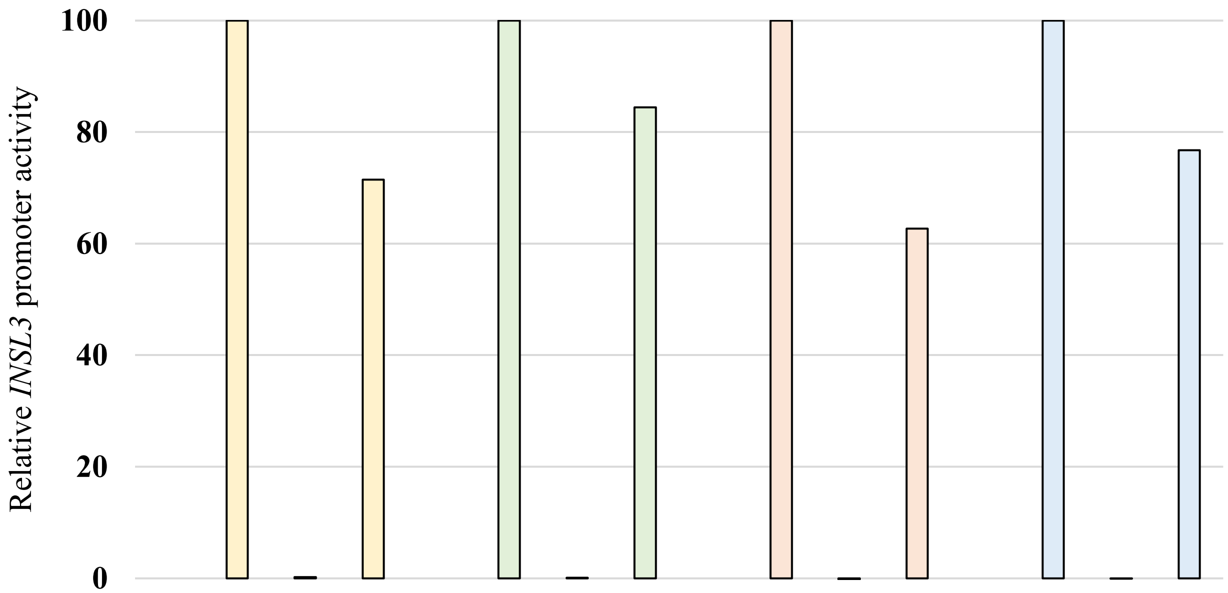


| | | | | | | | | | | | | | | | | |
|---|---|---|---|---|---|---|---|---|---|---|---|---|---|---|---|---|
| **p*INSL3* (ng)** | 50 | 50 | 50 | 50 | 50 | 50 | 50 | 50 | 100 | 100 | 100 | 100 | 100 | 100 | 100 | 100 |
| **NR5A1 (ng)** | - | 3.1 | - | 3.1 | - | 6.2 | - | 6.2 | - | 6.2 | - | 6.2 | - | 12.5 | - | 12.5 |
| **COUP-TFII-WT (ng)** | - | - | 3.1 | 3.1 | - | - | 6.2 | 6.2 | - | - | 6.2 | 6.2 | - | - | 12.5 | 12.5 |

Supplementary figure S3.
